# Comparative Evaluation of Three Calcium Silicate-Based Materials for Direct Pulp Capping: An *In Vivo* Study in Mice

**DOI:** 10.1101/2025.05.29.656939

**Authors:** Fatma Fenesha, Aonjittra Phanrungsuwan, Dane Kanniard, Brian L. Foster, Anibal Diogenes, Sarah B. Peters

## Abstract

**Introduction:** Vital pulp treatments (VPT) aim to conservatively manage deep caries and/or damage to preserve the vitality and function of the pulp. This minimally invasive approach is still debated when performing direct pulp capping (DPC), a treatment in which a protective biocompatible material is directly placed over the dental pulp, due to microbial risks inherent with pulp exposure and difficulties in sealing the exposure to protect the pulp from subsequent bacterial ingress. Several limitations associated with Mineral Trioxide Aggregate (MTA) such as long setting time and discoloration have promoted the development of next-generation MTA derivatives with enhanced physical, chemical, and biological properties. As these materials are relatively new, existing studies are limited in scope, lacking a comprehensive assessment of both reparative dentin formation and the sealing ability, which are critical parameters for determining long-term clinical success. These assessments require preclinical models, and while the mouse model offers the opportunity to explore the molecular mechanisms guiding reparative dentinogenesis, the materials and techniques optimized for human dentition present technical challenges in mice due to their small size.

**Aim:** We aimed to optimize a DPC technique to use in mice by comparing the tissue responses and the sealing ability of three calcium silicate capping agents: Bioceramic putty (BC), Biodentine (BD), and Theracal-LC (TLC) to a control material known to seal well but cause desiccation that would lead to tissue damage; Cavit-G (CG).

**Material and methods:** DPC was performed on the maxillary first molar of C57BL/6 female mice. Reparative dentin was assessed with micro-computed tomography (micro-CT) and histological assessment of dentin bridge formation at 21 days. The sealing ability of the capping materials was evaluated using micro-CT.

**Results:** All three calcium silicate materials showed good biocompatibility and the ability to form a dentin bridge. Micro-CT quantifications of material voids demonstrated superior seals with bioceramic putty and Theracal-LC groups as compared to Biodentine.

**Conclusions:** Ready-to-use, premixed capping agents exhibit better sealing ability in mice, while promoting dentin bridge formation to protect the dental pulp.

## Introduction

Dental caries (tooth decay) is one of the most common chronic diseases worldwide [1, 2], affecting people of all ages with a prevalence of 46.2% in primary teeth and 53.8% in permanent teeth [1–3]. The dental pulp is the innermost part of the tooth, a soft connective tissue that consists of nerves, blood vessels, and a wide array of cells. It plays a critical role in tooth formation, sensation, nutrition, and repair, and it ensures the long-term health of the tooth [4]. When the pulp is healthy, it keeps the tooth alive and functional. However, if the pulp becomes damaged or infected due to deep caries, trauma, or other factors, it can lead to pulp inflammation or pulpitis. Management of deep caries lesions, in which the infection extends close to or deep into the pulp, depends on the stage of inflammation (reversible or irreversible pulpitis) and requires careful planning to preserve tooth vitality and prevent the recurrence of the infection [5–8]. Currently, the most widely used treatment modality for pulpitis in adults is root canal therapy (RCT), which involves completely removing and replacing the dental pulp with a root canal sealing material. While this method has proven effective in addressing pulp inflammation and saving the tooth, it comes with certain limitations. It may render the tooth brittle in the absence of its natural pulp and blood supply, both of which are crucial for maintaining vitality and flexibility. Furthermore, if the root canal is not properly sealed or if the restoration leaks, bacteria can re-enter the tooth, causing reinfection, leading to apical periodontitis or abscess formation [9, 10]. Due to the growing recognition of the crucial function of dental pulp, modern dentistry is increasingly focusing on preserving the vitality of this tissue, reflecting a shift from conventional RCT to less invasive dental procedures such as VPT [11–13].

The American Association of Endodontists defines VPT as ‘’a treatment that preserves and maintains pulp tissue that has been compromised by trauma, caries or restorative procedures.’’ [14]. In this sense, VPT can involve directly (DPC) or indirectly capping the dental pulp (IPC), and full or partial pulpotomy [7, 15]. The success of VPT depends on the precise diagnosis and stage of inflammation, as well as selection of the pulp capping agent [16, 17]. The ideal pulp capping material should facilitate the formation of reparative dentin and the preservation of a vital dental pulp [18]. Several pulp capping agents have been introduced into the market **(Table 1)**, offering various success rates, advantages, and disadvantages. While some materials, like calcium hydroxide (CH), are widely recognized for their affordability and long track record [19, 20], they lack long-term durability and ability to adhere to the dentin surface. In addition, CH has been widely criticized for the tunnel defect-like phenomena in the formed reparative dentin [21–23], potentially leading to microleakage and bacterial invasion into the dental pulp [24]. To overcome the drawbacks of CH, calcium silicate cement-based materials were introduced such as MTA and MTA-derived materials. MTA was introduced as a therapeutic bioactive material that quickly gained popularity due to its’ biocompatibility, sealing ability, and excellent clinical success [25–27]. However, MTA has several limitations such as high cost, long setting time, tooth discoloration and handling challenges. Based on this, a search for a newer generation of endodontic materials, introduced some enhanced derivatives of calcium silicate capping agents. Biodentine (Septodeont, Saint-Maur-des-Fosses, France), was developed as a dentin substitute [28–30] and demonstrated success rates comparable to MTA in both short and long-term follow-up visits [19, 25, 31–33], but with Biodentine exhibiting better handling properties and quicker setting time than MTA [34]. However, both MTA and Biodentine are supplied in powder/liquid form, which requires a separate mixing step before application. This could lead to inconsistent chemical and mechanical properties of the capping agent due to variabilities in the powder-to-liquid mixing ratios [35, 36].

**Table 1:** Summary of common pulp capping materials:

| Material | Composition | Advantages | Disadvantages | Setting type/time |
| --- | --- | --- | --- | --- |
| Dycal<br>Pulpdent<br>Pulpcal | Base: Zinc oxide, calcium phosphate, calcium tungstate<br>Catalyst: Calcium hydroxide, zinc oxide and titanium dioxide. | Easy to work with, sets quickly.<br>Cost-effective [20]. | Limited antibacterial activity over time [60]<br>Poor cohesive strength [61].<br>Greater solubility, and marginal leakage [62, 63].<br>Tunnel defect in formed reparative dentin [21, 22]. | Self-curing<br>2-3 min |
| Cavit-G<br>Zinogen | Zinc oxide eugenol | Reduces inflammation [64]. | Releases eugenol even in low concentrations can be cytotoxic to dental pulp cells [65, 66] | Self-curing<br>5 –10 min setting continues for 1-2 hours |
| GIC<br>Fuji | Glass ionomer/<br>Resin modified glass ionomer | Excellent bacterial seal [67]<br>Fluoride release [68, 69].<br>Bond to both enamel and dentin.<br>Good biocompatibility [70]. | Cytotoxic when in direct cell contact and lack of dentin bridge formation (41) [71]. | Self-curing<br>3-5 min |
| Mineral trioxide aggregate (MTA) | Tricalcium and dicalcium silicate, tricalcium aluminate, tricalcium and silicate oxides and bismuth oxide. | Good biocompatibility [7, 27] More predictable hard tissue barrier formation than CH [72].<br>Antibacterial properties [73].<br>Osteoinductive [74]. | Poor handling characteristics [75].<br>Long setting time [76]. | Self-curing<br>30 min – 2 hours |
| Biodentine | Powder: tricalcium silicate, calcium carbonate and zirconium oxide.<br>Liquid: water and calcium chloride. | Biocompatible [77].<br>Bond to deep moist dentin [32, 78]<br>Induces mineralized bridge formation [45].<br>Faster setting time and good physiochemical properties [28]. | Shorter working time and limited radiopacity [79].<br>Difficulty in achieving the desired consistency.<br>Poor bonding with overlying restorations [80] | Self-curing<br>12 min |
| Theracal LC | Calcium silicate, barium sulphate, barium zirconate, fumed silica, and Bis-GMA and PEGDMA resin. | Low solubility and good mechanical properties [81].<br>Easy handlings, convenient and short setting time [82].<br>Induced dentin bridge formation [83]. | Long-term efficacy is limited [84].<br>Higher cytotoxic effects and reduced cell viability [77].<br>Reduced radiopacity | Light-cured<br>10 sec |
| Premixed Bioceramics<br>(EndoSequence BC, TotalFill BC RRM, iRoot BP Plus) | Calcium silicate, zirconium oxide, calcium phosphate, calcium sulphate and fillers. | Superior handling, uniform capping consistency and convenient manipulation [85].<br>Biocompatible [77] and ability to induce mineralization and odontoblast differentiation [40]. | Limited long-term data [42].<br>Tooth discoloration [44]. | Self-curing<br>15 -20 min |

To further improve handling and maneuverability in clinical settings, ready-to-use capping materials were developed such as Theracal LC (Bisco, Scaunburg, IL, USA) and premixed bioceramics such as iRoot BP Plus (Innovative Biocermaics, Vancouver, BC, Canada), TotalFill RRM putty (FKG, La-Chaux-de-Fonds, Switzerland) and EndoSequence BC putty (Brasseler, USA, Savannah, GA). Theracal LC is a hybrid, light-cured resin-modified calcium silicate material. The addition of the resin monomer provided some advantages such as enhanced physical properties, shorter setting time, and user-friendliness. Despite its advantages, there are concerns regarding the potential cytotoxic effects from residual resin monomers coming in contact with the dental pulp tissue [37–39]. Regardless, premixed bioceramics have been shown to possess more favourable biocompatible properties in inducing reparative dentin formation due to their apatite-forming ability [40, 41]. Additional advantages have also been reported, including a uniform consistency, minimum technique sensitivity [42], antibacterial properties [43], and versatile applications [44]. These ready-made materials have been mostly tested as endodontic sealers and root canal obturation agents, and few studies have been conducted to evaluate these materials as pulp capping agents [42]. Furthermore, most studies have focused primarily on reparative dentin formation as the main outcome when evaluating calcium silicate-based capping agents [27, 31, 41, 45, 46] and have not assessed critical factors such as sealing efficiency and microleakage *in vivo* [47–49]. This highlights a gap in understanding the biocompatibility and mechanical performance of currently available pulp capping agents and emphasizes the need to perform comprehensive analyses in preclinical models *in vivo*.

It is crucial to note that while human studies provide important evidence on materials performance, the analyses rely on two-dimensional radiographs, which lack the resolution to detect thin or early dentin bridge formation (<0.5 mm) [50]. Preclinical models allow for a multidimensional assessment of the healing response, such as the structural features of the dentin bridge. Rats are among the most widely used preclinical models for endodontic research due to their size, cost-effectiveness, and relatively straightforward handling in laboratory settings [51–58]. However, rats do not provide the genetic flexibility that mice do to investigate the underlying cellular and molecular mechanisms driving tissue repair [59]. Despite these advantages in mice, their small mouth and tooth size make these procedures technically challenging, but surgical operating microscopes have improved the ability to perform endodontic and dental procedures in smaller animals. Therefore, the aim of this study is to refine a pulp capping technique in mice by testing three calcium silicate capping materials: Biodentine (BD), Bioceramic putty (BC) Putty, and Theracal LC (TLC). We assessed the materials’ effects on reparative dentin formation (via microCT and histology), and sealing quality (via void analysis). We hypothesized that comparing these materials head-to-head in a controlled mouse model would create conclusive evidence regarding their impact on dentin healing and better inform material selection and future preclinical and translational research in regenerative endodontics.

## Materials and methods

This study incorporated the Preferred Reporting Items for Animal studies in Endodontology (PRIASE) 2021 guidelines [86, 87].

### Animals

All mouse experiments were approved by the Institutional Animal Care and Use Committee (Protocol #2020A00000023-R1). Three-month-old wild-type (WT) C57BL/6 female mice weighing from 17 – 22g were used. To ensure ethical and responsible conduct of animal research, the number of mice in this study was minimized in accordance with the ARRIVE guidelines [88]. The mice were maintained by qualified caretakers in a sterile setting with controlled humidity, temperature, and lighting. We conducted the study using only female mice to minimize the gender effects on dentin healing.

### Direct pulp capping procedure

A total of 16 mice were randomly divided into 4 experimental groups (n=3-4 mice/group); **CG**, Cavit-^i^G (3M ESPE, St Paul, MN) with a collagen sponge (Zimmer Plug^®^ and Optimaix 3D^®^), **TLC**, Theracal LC (Bisco, Schaumburg, IL, USA), **BC**, Bioceramic RRM putty (Brasseler USA, Savannah, GA), or **BD**, Biodentine (Septodont, Saint-Maur-des-Fossés, France). The mice were anesthetized with isoflurane and weighed before the experiment. We used a standardized mouse tooth injury model as described by Yang et al [89] in **Figure 1A**. A type ll or intermediate injury was performed involving the deep dentin with partial loss of pulpal tissue stimulating the formation of reparative dentin as part of the healing response [90]. The maxillary first molar was first disinfected with 70% ethanol and a cavity was drilled without exposure using a 0.3 mm round carbide bur (Komet, USA) on the mesial surface of both maxillary first molars with constant cooling. The mesial horn of the dental pulp was then exposed using #15 K endodontic file (Dentsply, Switzerland) with a pinpoint bleeding confirming the exposure, after which haemostasis was achieved using a sterile paper point. The exposure was capped with one of the four capping materials; CG, BC, BD or TLC, and then sealed with glass ionomer cement (Fuji IX, GC) as per the manufacture instructions **(Figure 1B).** Each capping material was applied in a single layer and allowed to set according to the manufacture instructions **(Table 1).** Excess material was removed, and the tooth and gingival tissues were cleaned with a cotton tip soaked in sterile distilled water. The mice were returned to their cages with no food for 30 minutes and then supplied with soft diet gel (W.F.Fisher and Son, USA) for 24 hours to minimize discomfort during mastication. The mice were maintained on standard chow and monitored for changes in eating behaviours. Weights were recorded at day 0- and 21-days post capping.

**Figure 1:**
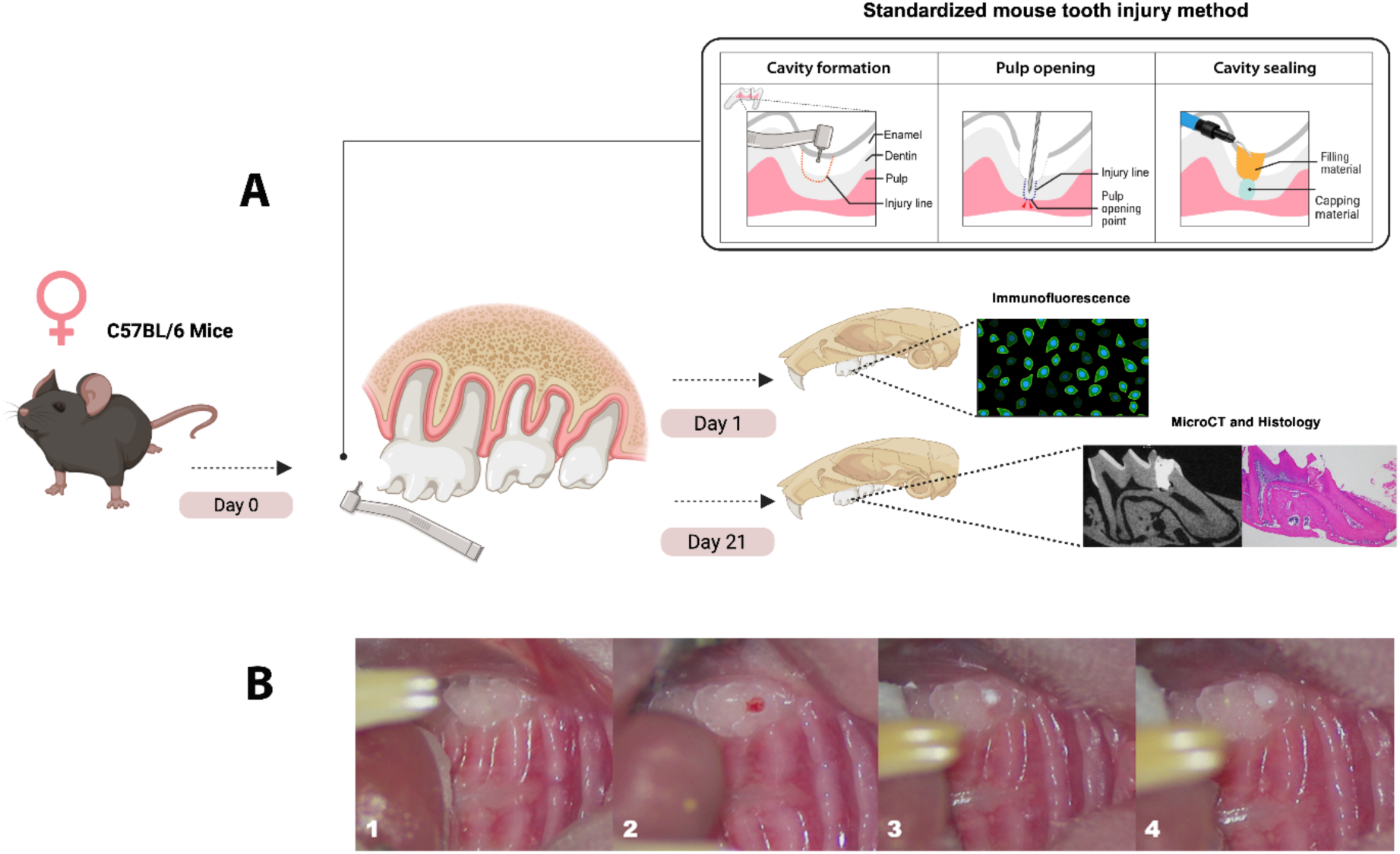
Experimental design and standardized pulp capping procedure: **(A)** Schematic representation of the experimental workflow and pulp capping technique using a standardized mouse tooth injury method. Maxillae samples were collected at two time points: Day 1 for immunofluorescence labelling of immune cells, and Day 21 for Micro-CT and histological analysis. **(B)** Sequential images of the pulp capping procedure. (1) Unprepared maxillary first molar in a mouse. (2) Cavity preparation using a 0.3 mm round bur, followed by pulp exposure with a #15K endodontic file. (3) Application of the pulp capping material over the exposed pulp tissue. (4) Placement of a glass ionomer restoration to seal the cavity. Standardized mouse tooth injury illustration adopted from <u>Yang *et al* 2023.</u> Illustration created with Biorender.

### Micro-computed tomography (µCT)

The mice were sacrificed at 21 days post-capping using intracardiac perfusion buffered with 0.5M PB with 4% paraformaldehyde. Hemimaxillae were fixed in 4% paraformaldehyde for 2 hours at room temperature, followed by two rinses with PBS and transfer to 70% ethanol for micro-CT scanning (Scanco Medical, Brüttisellen, Switzerland) at 70 Kvp, 6 µm voxel dimension, 0.5 Al filter, and 1,200 ms integration time. Known density hydroxyapatite (HA) phantoms (0, 100, 200, 400, and 800 mg HA/cm^3^) were scanned at the same parameters and used to quantify a standard linear curve to formulate the density unit of each sample as mentioned in our previous article [91]. DICOM reconstructed files were analyzed with AnalyzePro 15.0 (AnalyzeDirect). The first maxillary molar and associated alveolar bones were segmented as regions of interest (ROI). The ROI was outlined as the area between 240 µm mesial from the most mesial point of the mesial maxillary first molar root and 240 µm distal from the most distal point of the distal maxillary molar root. Reparative dentin was manually segmented at 240-1,600 mg HA/cm^3^. Periodontal ligament (PDL) space of the mesial maxillary molar was manually segmented using the apical 1/3 of the total mesial root length. The sealing ability of the four capping materials was assessed by measuring the voids percentage in the capping materials. The capping material was outlined, and the voids were manually segmented at 2300-2811 mg HA/cm^3^. This range was selected to effectively exclude enamel from our analysis, as enamel possesses a significantly higher mineral density compared to dentin and other dental tissues. The total volume of each capping material and the internal void volume were quantified using high-resolution micro-CT. Segmentation was performed based on grayscale thresholding to differentiate between the capping material and voids. All measurements were conducted using consistent threshold values and image analysis parameters to ensure comparability across samples. The voids percentage of each capping material was calculated using the following equation: Voids percentage = (voids volume / total void + capping volume) X 100.

### Histological examination

The mice were sacrificed at 21 days post-capping using intracardiac perfusion buffered with 0.5M PB with 4% paraformaldehyde. Hemimaxillae were fixed in 4% paraformaldehyde for 2 hours at room temperature and decalcified in poly-Nocal (Polysciences) for 5-7 days. Tissues were then dehydrated and embedded in paraffin for 6 µm sections along the midsagittal axis. The sections were stained using Hematoxylin and eosin (H&E) and Masson’s Trichrome stains according to standard protocols and mounted with a mounting solution.

### Statistical analysis

Mean ± standard deviation values were shown in the graphs. The normality test was achieved using Shapiro-Wilk test. Data were calculated with a one-way analysis of variance (ANOVA) test and Tukey’s multiple comparisons test to compare between the capping materials, where p = 0.05 or less was considered statistically significant. Data analysis was performed using statistical software (Prism 10.4.1; GraphPad Software). Sample size calculation was based on previous studies evaluating comparable outcomes [31, 41], with a significance level (α) of 0.05, and a desired power (1-β) of 0.8, the analysis indicated a total number of 16 mice (n=3-4 mice/group) with bilateral molars to be sufficient to detect statistically significant differences.

## Results

No adverse events occurred, nor any mice were lost during the experiments. A total of 16 mice (32 molars) were included in the final analysis.

### Differential dentin bridge formation induced by calcium silicate-based pulp capping agents

Micro-CT analysis was employed to assess the formation of tertiary dentin and the morphology of PDL space (**Figure 2A-H and 2L**). At 21 days, we found that the CG group exhibited a statistically significant reduction in tertiary dentin formation compared to the other groups. Ectopic calcifications were observed in the CG cohort (see **Figures 2A and 2I**). A notable disparity in tertiary dentin volume was identified between the TLC group and the BD treated group (**Figure 2I**). No significant differences were detected in the density of tertiary dentin between experimental groups (**Figure 2J**). The analysis of the PDL space surrounding the apical one-third of the mesial roots was conducted to evaluate the inflammatory response elicited by each capping material. Interestingly, the CG restorations led to enlarged PDL space relative to the BC, BD, and untreated group. However, we did not find any significant difference between the untreated group and TLC, BC, and BD groups. This observation suggests that CG treatment induces pulpal inflammation (**Figure 2K-L**) and **(Figure 3A-D).**

**Figure 2:**
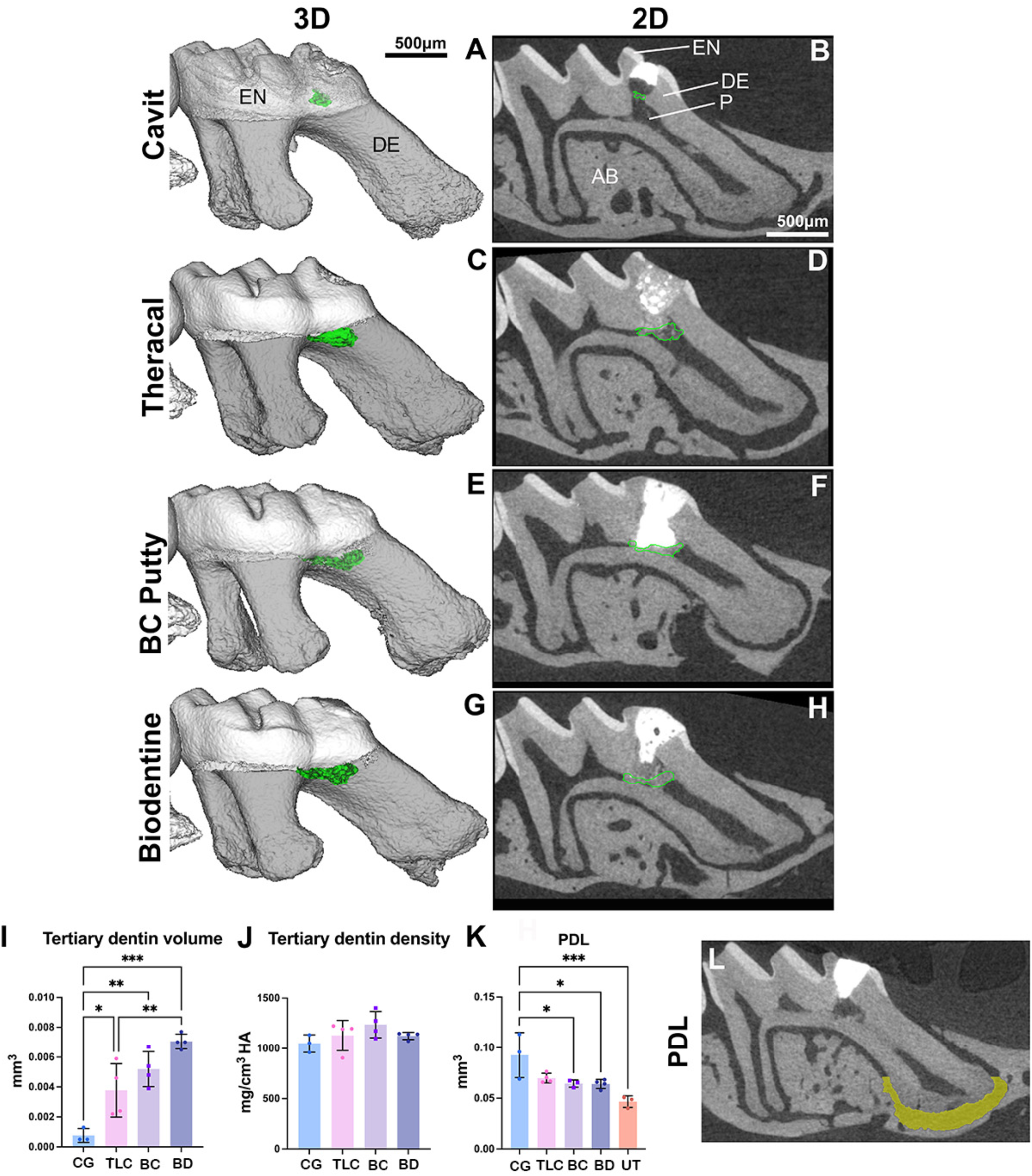
Micro-CT analysis of reparative dentin formation and PDL widening at 21 days post-capping: (A,B) Cavit-G group (negative control) displaying signs of random opacities along the root canal. **(C-H)** representative 3D and 2D slices demonstrating reparative dentin (outlined in green) formed which appear denser and localized beneath the pulp capping material. Quantification of reparative dentin volume **(I)** and density **(J)** showing significant differences in pulp capping volume between each capping material; TLC, BC and BD with CG. No significant differences were observed in density between the four materials. Region of interest (ROI) for measuring PDL widening is represented in **L**. The PDL was manually segments at the apical 1/3 of the total mesial root length of the injured maxillary first molar. Quantification of the PDL widening presented in **K**, significant differences were observed between BD and CG, BC and CG, and UT (untreated) and CG. No significant differences were observed between the three capping materials (BC, TLC and BD) and UT group. Scalebars are 50μm in **B**. \**P* < 0.05, \*\**P* <0.01, *** *P* < 0.001 by one-way ANOVA. P: pulp, DE: dentin, EN: enamel, AB: alveolar bone, CG: Cavit-G, TLC: Theracal LC, BC: BC putty, BD: Biodentine.

**Figure 3:**
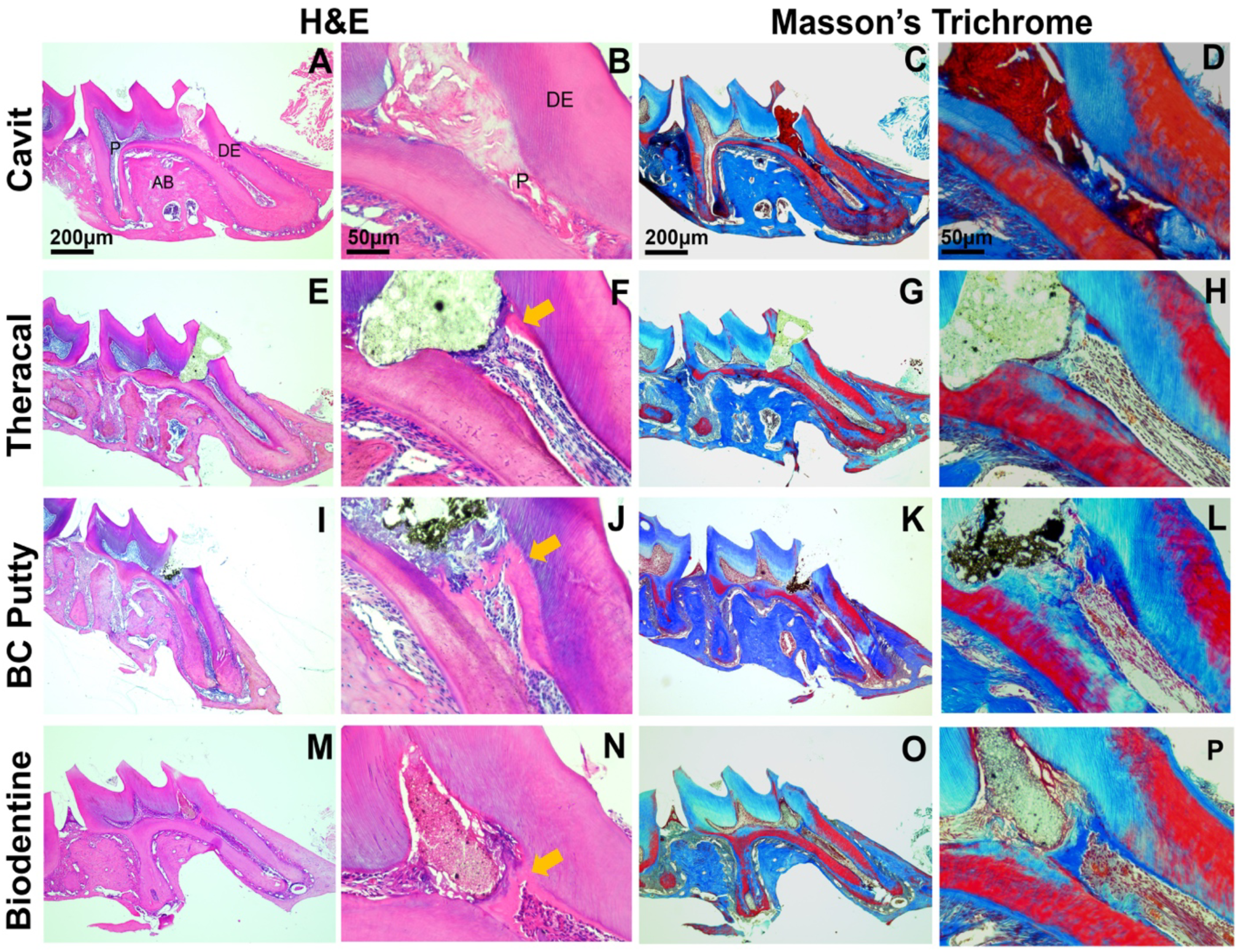
Histopathological evaluation at day 21 post-capping using Hematoxylin-eosin and Masson trichrome staining. **(A-D)** Cavit-G group (negative control) displaying no evidence of reparative dentin, absence of odontoblasts and dental pulp cells, and extension of the inflammation along the mesial root canal. **(E-H)** Theracal – LC group. **(I-L)** BC putty group. **(M-P)** Biodentine group. Images **E-P** demonstrating reparative dentin formation indicated by the yellow arow in **F**, **J** and **N**. Scalebars = 200μm in **A** and **C** and = 50μm in **B** and **D**. P: dental pulp, DE: dentin, AB: alveolar bone.

### Enhanced sealing ability with premixed calcium silicate-based pulp capping agents

DPC with BD resulted in the highest material voids (mean = 13.4 %), with TLC pulp capped molars demonstrating 8.31% average void space, and BC pulp capped molars demonstrating an average of 6.39%. Significant differences in void percentage were found between the BD and BC groups (*P* = 0.008). No significant differences were found between TLC and BC in void percentage (**Figure 4 A-M**). No significant differences were found in the total capping material volume between all the groups (*P* > 0.05). **Figure 4** shows representative images of the total capping and void volume in 2D slices and 3D segmentation.

**Figure 4:**
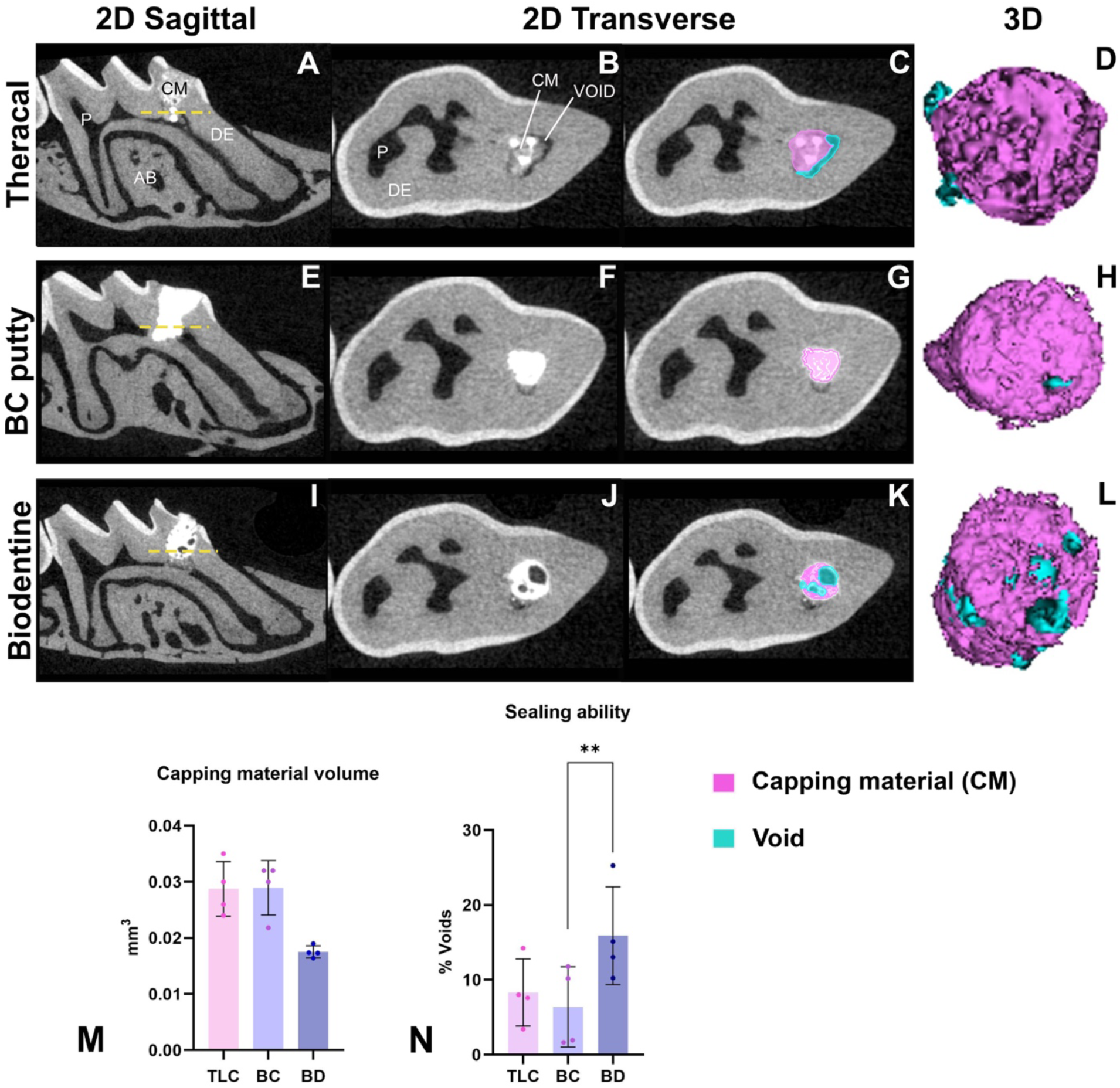
Micro-CT analysis of the sealing ability of the capping materials at 21 days post-capping. This illustration shows sagittal cross sections of the capped first maxillary molar presented in **A,E** and **I.** Trans axial 2D slices sectioned at the dashed yellow line in **(B-K)** displaying the total capping and void areas in TLC, BC and BD. 3D rendering of the three capping material (purple) and associated voids (blue). Quantification of the capping material volume **(M)** and voids % in **(N)** between the three capping materials (TLC, BC, and BD). Scalebars are 100μm. \**P* < 0.05, by one-way ANOVA. P: pulp, DE: dentin, AB: alveolar bone, CM: capping material, TLC: Theracal LC, BC: BC putty, BD: Biodentine.

## Discussion

In recent years, there has been a growing interest in VPT, as an alternative to RCT [92–96]. Traditionally, the focus of VPT was primarily aimed at maintaining the vitality of the radicular dental pulp after minor injuries or caries exposure. This was typically indicated for primary teeth and immature permanent teeth with deep caries to allow continued root formation and apex closure [97]. Today, vital pulp therapy has evolved with a broader scope, no longer restricted to immature teeth. It is now a viable treatment option for mature permanent teeth, supported by advancements in biomaterials and minimally invasive techniques [5]. Despite the potential of vital pulp therapies to enhance the pulp’s healing in older adults, their reported success rates vary widely from 42.9% to 81.5%, and generally lower compared to full pulpectomy [23, 98]. This unpredictability hinders the widespread clinical adoption and success of VPTs, demanding further research to improve clinical outcomes. In response, bioengineers have made significant advances in biomaterials to enhance the biocompatibility and sealing outcomes of their pulp capping materials. As a result, MTA has largely replaced calcium hydroxide as the gold standard for direct pulp capping, due to its superior sealing ability, biocompatibility, and long-term clinical outcomes [26]. Newer calcium silicate-based cements were then developed to address the drawbacks associated with MTA, including its’ long setting time, difficult handling properties, discoloration, and relatively high cost. Materials such as Biodentine, BC Putty, and Theracal LC offer improved clinical usability through faster setting times, premixed formulations, and enhanced mechanical properties, while retaining the biological benefits of MTA. Despite promising *in vitro* and clinical data, variability in composition, setting mechanisms, and long-term outcomes have prevented direct comparisons between these newer materials to make informed decisions on material selection. Further preclinical research is essential to understand how each material and technique impacts sealing ability, pulp inflammation, and pulp healing.

Although calcium silicate-based materials have shown promising results in preclinical and *in vitro* studies [19, 25, 31–33, 40, 41], translating these findings directly to clinical success in human teeth is not always straightforward. This is because studies are often conducted using different animal models (e.g., rats, dogs, or mice), which each have unique pulp anatomies, healing capacitates, and immune responses that differ significantly from human teeth [26, 59]. The choice of model depends on the specific research goals, ethical considerations, and resources available. This study focused on developing the techniques and materials in mice to allow for future studies utilizing transgenic mouse models with global and conditional deletion and/or overexpression of genes. Our study has established a standardized and reproducible technique for performing DPC in mice, enabling consistent assessment of pulp healing and material performance in a genetically tractable model.

In the present study, we focused on comparing three commercially available calcium silicate pulp capping agents to help guide evidence-based material selection and translate the findings into clinically reliable outcomes. Our study demonstrated that all three calcium-silicate-based materials induced reparative dentin formation by day 21, demonstrating their biocompatibility and capacity to support pulp healing. Significant differences in tertiary dentin volume were observed between the three calcium silicate materials and the Cavit-G control; however, no significant differences in dentin density were detected among any of the groups. Our findings are in alignment with previous studies evaluating calcium silicate-based materials as pulp capping agents [40, 45, 77, 83], further supporting their effectiveness in promoting reparative dentin formation and pulp healing.

A key factor for the success of VPT is achieving a well-adapted void free seal to prevent microleakage. An inadequate seal is a major cause of failure in endodontic procedures, as it allows bacterial penetration, leading to recurrent inflammation and compromised treatment outcomes [48, 49]. A material that promotes dentin formation but fails to provide an effective seal compromises pulp protection and thus would not be clinically viable for long-term performance. In this study, we evaluated the sealing ability of different capping materials by measuring the voids percentage. Biodentine exhibited higher voids percentage than BC putty and Theracal LC when combined with glass ionomer restorations, suggesting that the BC putty may offer superior sealing properties compared to the other materials, potentially reducing the risk of microleakage and associated complications. These findings are in agreement with prior studies assessing the sealing ability and microleakage of pulp capping materials [99–104]. In an *in vitro* study, Makkar et al evaluated the sealing ability of Biodonetine and Theracal LC against the conventional MTA using confocal laser microscopy and Rhodamine B fluorescent dye. Both Biodentine and MTA showed comparable micropermeability at the material dentin interface. However, Theracal LC exhibited significantly higher sealing ability than the other two materials [99]. One possible explanation for the lower sealing ability of Biodentine, is the variability introduced by its powder-to-liquid mixing process, which may lead to inconsistencies in material properties compared to ready-to-use materials. On the contrary, Anusha et al compared the microleakage of Biodentine with Theracal PT; a newer dual-cured calcium silicate capping agent. Their findings demonstrated no significant differences in microleakage scores between the two capping materials [100]. Nevertheless, it should be emphasized that TheraCal PT was developed to improve the cytocompatibility of resin-based calcium silicate capping agents by reducing the resin content, in contrast to its predecessor, TheraCal LC [101]. While its cytotoxicity has been shown to be comparable to Biodentine [102, 103], the reduced resin content in TheraCal PT could potentially reduce its mechanical strength and sealing efficacy [104].

To the best of our knowledge, the sealing ability of BC putty as pulp capping agent has not been previously evaluated. De-Deus et al assessed the interfacial adaption and internal gaps of Endosequence BC as a root canal sealer compared with the epoxy resin-based AH Plus sealer. Although both groups demonstrated gaps at the interface, AH plus displayed better sealing adaptability than BC sealer [105]. Raju et al compared the apical sealing ability of ProRoot MTA, Biodentine and Total Fill BC as root end filling materials. TotalFill BC RRM Putty exhibited minimal microleakage along the root canal walls *in vitro* [106]. The literature remains inconsistent regarding the sealing ability of endodontic materials, attributed to the methodological differences across studies. In our study, we provide a novel method with which to quantify sealing ability and emphasize its ease-of-use with preclinical rodent models for the community of researchers involved in improving VPTs.

This present study contributes several data points to the growing body of evidence on calcium silicate-based materials as pulp capping agents. First, the study employed a standardized and reproducible pulp capping technique, ensuring uniform application of materials and minimizing procedural variability. All procedures were performed by a single trained investigator to ensure consistency and minimize inter-operator variability. Second, the combined use of micro-CT imaging and histological analysis for the quantification of reparative dentin volume, along with the sealing ability, offers a more nuanced understanding of material performance than studies relying on a single outcome measure. Thirdly, assessing the sealing efficacy of a pulp capping material can help predict the long-term success and durability of the VPT. Lastly, comparing three widely used calcium silicate materials head-to-head under identical experimental conditions, can be used to inform both preclinical researchers and clinicians on materials to utilize in their studies and practice.

This in vivo study was conducted in response to the inconsistency and lack of standardization in previous VPT research, particularly the use of varying animal models. Moreover, the sealing ability of these newer materials remains underexplored. Therefore, a standardized DPC mouse model was necessary to enable direct comparison. Nonetheless, this study has certain limitations. One limitation of this study is the use of sound healthy molar teeth rather than a pulpitis model. While mechanically exposed healthy pulps allow for controlled comparisons of reparative dentinogenesis and inflammation, they do not fully replicate the clinical environment where pulp capping is typically indicated. As such, the healing dynamics observed in this model may differ from those in a diseased state, potentially affecting the translational relevance of the findings. Additionally, while micro-CT provides a valuable volumetric assessment of voids and material placement, it does not capture the microscopic interaction between the capping material and dentinal surfaces [107, 108]. This includes adhesion at the dentin interface and penetration into dentinal tubules, which are critical for long-term sealing efficacy. Future studies should incorporate advanced techniques to better characterize capping material adaptation and bonding at the dentin interface and with the overlaying restoration.

## Conclusions

In this study, all three calcium-silicate-based pulp capping materials; Biodentine, BC Putty, and TheraCal LC successfully induced reparative dentin formation, demonstrating their biocompatibility and healing potential when compared to the negative control; Cavit-G. Among them, BC Putty and Theracal LC exhibited superior sealing ability, characterized by a greater extent of void-free areas, suggesting enhanced marginal adaptation. These findings highlight the importance of evaluating both biological and physical properties to inform the selection of pulp capping materials in clinical practice.

